# Multisensory integration in Peripersonal Space indexes consciousness states in sleep and disorders of consciousness

**DOI:** 10.1101/2024.10.25.619776

**Authors:** Tommaso Bertoni, Giulia Ricci, Jane Jöhr, Brunella Donno, Jacinthe Cataldi, Julia Fellrath, Aurelie Stephan, Carolina Foglia, Sandro Lecci, Florian Dauvin, Marina Lopez Da Silva, Mattia Galigani, Polona Pozeg, Vincent Dunet, Marzia De Lucia, Jean-Paul Noel, Elisa Magosso, Karin Diserens, Francesca Siclari, Andrea Serino

## Abstract

Conscious experience encompasses not only the awareness of external objects, but also a phenomenal representation of the embodied subject of the experience. The latter is mediated by the integration of multisensory stimuli between the body and the environment, a process mediated by the Peripersonal Space (PPS) system. Here we thus tested the hypothesis that a neural marker of PPS representation may index the presence of conscious experience. Using high-density EEG in awake participants, we identified a “PPS index”, characterized by high-beta oscillations in centroparietal regions during the integration of audiotactile stimuli presented near versus far from the body. We then examined this marker across two models of altered consciousness, i.e., sleep and disorders of consciousness. The PPS index persisted during dreaming and waking conscious states but was absent during dreamless, unconscious states. Moreover, the same index predicted behavioural measures of consciousness and clinical outcome in patients recovering from disorders of consciousness. These results suggest that multisensory integration within the PPS is tightly linked to the presence of conscious experience.

## Introduction

The neural correlates of consciousness have sparked a vibrant debate for more than 30 years ^1–3^. The majority of theories ^4^ focus on the mechanisms underlying the awareness an external stimulus (e.g., visual in the recent major initiative from the Cogitate Consortium ^5^). Neurophenomenological approaches highlight the investigation of the structure of consciousness, in addition to its content ^6,7^. In this framework, subjective experience implies a phenomenal representation not only of the object, but also of the subject of experience, “here and now” within one’s environment ^8,9^. Such representation, also termed minimal selfhood ^8^, is thought to emerge from the continuous and coherent integration of multisensory information from one’s own body and the environment, leading to the concepts of Bodily Self-Consciousness and Bodily Self ^10–12^.

A neural system integrating bodily cues with external stimuli, considered to be key for the emergence of the Bodily Self, is the Peripersonal Space system (PPS). PPS is constituted by a network of neurons, primarily located in the posterior parietal cortex, premotor cortex, temporo-parietal junction, and putamen, that integrates tactile information on the body with external - visual or auditory - stimuli as a function of their distance, in a body-centered reference frame ^13,14^. Through this process, PPS builds a representation of the body in a space of potential interactions, which constitutes a basic interface between the self and the environment. Neurophysiological ^15^, behavioural ^16^, computational ^17^ and neuroimaging ^18,19^ evidence supports the view that PPS is involved in integrating multisensory stimuli underlying Bodily Self-Consciousness. Classic multisensory manipulations used to study Bodily Self-Consciousness, such as the rubber hand illusion ^20^, the full body illusion ^21,22^ or the enfacement illusion ^23,24^, tap into the spatio-temporal coherence of multisensory stimuli that PPS neurons process ^14,25,26^. Accordingly, studies have shown that PPS representation varies following changes in Bodily Self-Consciousness ^18,27,28^. Finally, our previous work demonstrated that, in the acute phase of patients with severe brain injury, the presence of an intact a PPS representation distinguishes cases suffering cognitive motor dissociation (CMD) within the whole population of patients with a disorder of consciousness (DoC). CMD patients exhibit a behavioural phenotype of DoC, but actually suffer from a motor efferent disorder limiting their ability to produce voluntary responses, despite being conscious and aware of their surroundings ^29, 30^.

Considering these converging findings about the role of PPS in self-consciousness, here we propose that the presence of a neural signature of PPS representation can be taken as a measure of minimal selfhood. In this perspective, a marker of PPS processing should be present during conscious states and vanish in the absence of consciousness. To test this hypothesis, we studied two models of changes in consciousness states, i.e., (1) wake and sleep cycles, and (2) patients recovering after pathological awakening from coma, exhibiting a DoC behavioural phenotype. We developed a neurophysiological marker of PPS processing, derived from intracranial and scalp electroencephalography (EEG) recordings in healthy participants ^31,32^, neurological patients ^33,34^ and newborns ^35^. Participants were presented with either unisensory or multisensory tactile and auditory stimuli, the latter occurring either near or far from the body (Fig. 1). Due to the neurophysiology of PPS neurons ^8^, under normal conditions, multisensory stimuli elicit a stronger response specifically when occurring near the body, and this is taken as a marker of PPS representation ^34^. We measured this marker in awake healthy individuals and contrasted it as a function of the level of consciousness across different sleep stages. Sleep is “an invaluable windows on consciousness” ^36^, as during sleep, the level of consciousness varies considerably, ranging from unconsciousness in dreamless sleep (no experience) to vivid conscious experiences in the form of dreams ^37,38^. While by definition, sleep involves a disconnection from the environment characterized by reduced responsiveness to sensory stimuli, studies show that the level of this disconnection may vary, and that windows of preserved behavioural responsiveness and sensory processing may exist during full-fledged sleep ^39–42^. We predicted that, if PPS processing captures a minimal form of selfhood characterizing consciousness, a marker of PPS representation should be found during sleep periods with consciousness - dreaming sleep - and not during sleep without consciousness - dreamless sleep. We then reasoned that if PPS processing is indicative of consciousness during sleep, the PPS marker may also index consciousness levels in patients with DoC phenotype and predict clinical recovery ^43^. We then studied their anatomical and functional neuroimaging data to investigate the neural structures related to our PPS marker.

**Figure 1.**
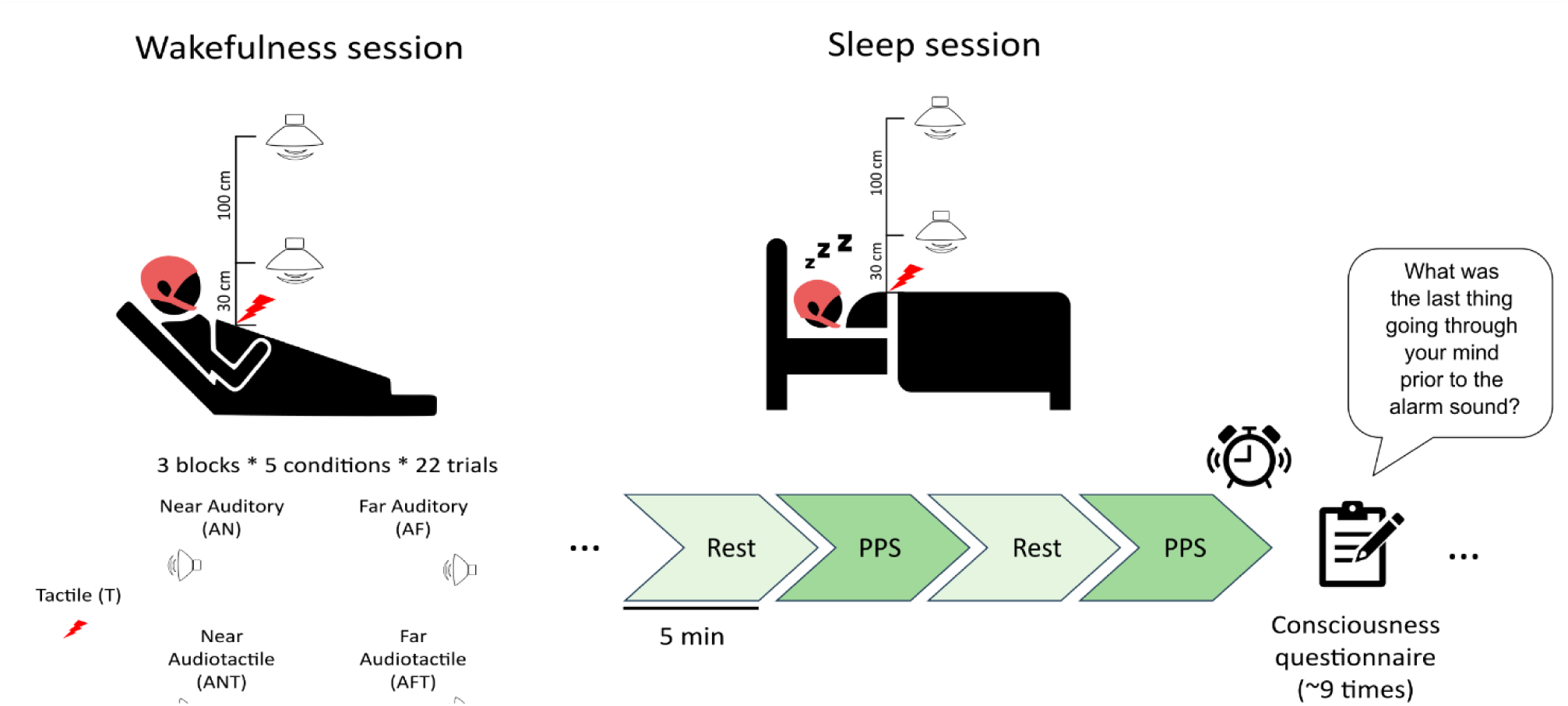
Sleep experiment design. The 5 experimental conditions are shown on the left. In the wakefulness session, three blocks of 110 stimuli (22 per condition, randomized) were administered (PPS paradigm). In the sleep session, participants received 5 minutes blocks with stimuli from the PPS paradigm as in the wakefulness session, alternated with 5 minutes blocks of resting state. About 9 times per night, they were awakened and responded a questionnaire about the presence and the content of conscious experience before the awakening. 256 channels EEG was recorded throughout the experiment.

## Results

### High beta oscillations index PPS representation during wakefulness

We first aimed at establishing a neurophysiological index of PPS representation which can be meaningfully compared across consciousness states. As an index of PPS representation, we searched for a space dependent multisensory effect, whereby stimuli presented at a given distance from the body (near vs. far) differently affect neural processing in unisensory (auditory) versus multisensory (audio-tactile) conditions. As in previous studies ^33,35^, we defined a PPS index as the near-far difference in power for unisensory inputs minus the near-far difference for multisensory inputs, applying the formula to spectral power rather than to ERP or GFP amplitude (see Methods). Statistically, this is equivalent to the interaction between distance (near-far) and modality of the presented stimuli (uni- or multi-sensory).

We leveraged such index to detect channels and frequency bands showing signatures of PPS representation in healthy awake participants. Hence, we first averaged the power spectra for each channel across five frequency bands (theta, alpha, beta low, beta high and gamma, see Methods) during the 1-second following stimulus onset. Then, we computed the PPS index for each channel and tested it against zero. We found one significant cluster in centro-parietal electrodes (p = 0.0084, see Methods) indicating PPS processing in the high beta (20-30 Hz) range (Fig. 2a). We then studied the frequency of the PPS effect in detail within this cluster, by computing the PPS index for all frequencies. The PPS index was significantly different from 0 (one sample t-test) between 22 and 37 Hz, peaking at 26 Hz (Fig. 2b), covering most of the high beta band. We then projected the high-beta PPS index to cortical locations following EEG source reconstruction (see Methods). We found significant activations in the bilateral precuneus, cingulate, intraparietal sulcus (IPS), in the left superior parietal lobule (SPL) and in the right inferior parietal lobule (IPL). We found further activations in the left somatosensory and motor cortex and small clusters in the superior temporal sulcus (STS), temporo-parietal junction (TPJ) and superior frontal gyrus (Fig. 2c). These activations largely matched PPS-related regions as identified by previous neuroimaging studies ^19^.

**Figure 2.**
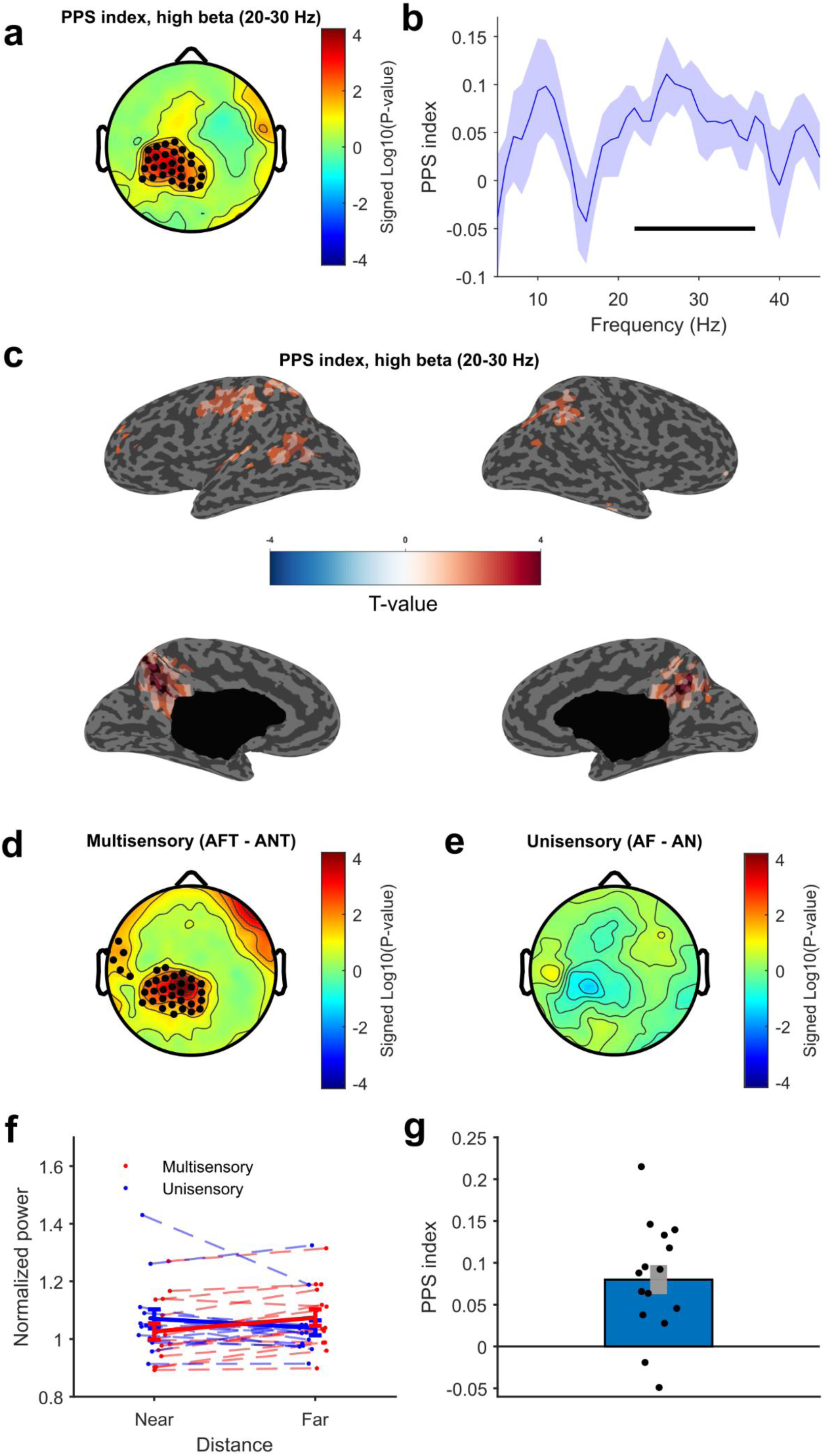
Spectral index of PPS representation during wakefulness. Panel (a) shows the topography of the PPS index in the high beta (20-30 Hz) frequency range. The black dots in the centro-parietal area denote electrodes belonging to the significant cluster. Panel (b) shows the PPS index by frequency in the significant cluster. Shades indicate standard errors, and the horizontal line indicates frequencies where the PPS index is significantly greater than zero. Panel (c) shows the cortical reconstruction of the PPS index, thresholded at p < 0.05. Panel (d) shows the comparison between near and far multisensory stimuli (AFT-ANT). Black dots denote electrodes belonging to the significant cluster. Panel (e) shows the topography of the near-far comparison within unisensory stimuli (AF-AN). Panel (f) shows power in the high beta band within the cluster identified in (a) in the four experimental conditions for each participant. Thicker lines represent means and error bars standard errors. Panel (g) shows the PPS index in the high-beta cluster for each participant (the error bar represents the standard error).

To further characterize this effect, we compared the response to multisensory (ANT vs. AFT) and unisensory (AN vs. AF) stimuli as a function of distance. We found a significant cluster, largely overlapping with the one found for the PPS index, when comparing multisensory stimuli (p = 0.0088; Fig. 2d). No significant cluster of electrodes emerged for unisensory stimuli (Fig. 2e).

A reduction in power in the alpha and beta bands typically indicates higher activation of the sensorimotor system ^44^, and a recent study showed suppressed beta activity in response to tactile stimuli coupled with looming visual stimuli ^45^. Coherently with these findings, post-hoc analyses within the cluster of significant interaction showed that near audiotactile stimuli induced a stronger high-beta desynchronization (power reduction) compared to far audiotactile stimuli (p < 0.001; Fig. 2f). No significant difference was observed for unisensory stimuli (p = 0.14). Thus, the PPS effect shown in Figure 2a was driven by a space-dependent modulation of multisensory processing. Figure 2g displays the high-beta PPS index in the cluster for all participants. The index was positive for all but 2 of the 15 participants.

### Spectral markers of PPS representation characterize conscious experience during sleep

We then studied the spectral markers of PPS representation previously identified during wakefulness across non-REM (stages N2 and N3) and REM sleep stages. Sleep is associated with broad fluctuations in the presence and content of conscious experience, spanning from absence of experience to vivid dreams. Although dreams were initially thought to occur almost exclusively during REM sleep ^46^, it is now well established that they are not exclusive to this stage ^38^, and recent studies have shown that they share similar cortical/EEG correlates across REM and NREM (N2 and N3) sleep ^37^. Thus, by using a serial awakening protocol ^47^, we defined dreaming and no-dreaming periods based on subjective reports. Following our hypothesis that PPS representation is a marker of minimal selfhood characterizing conscious experience, the high-beta PPS index found during wakefulness should be present during dreaming experience, and disappear during dreamless sleep, i.e., when no conscious content was reported.

First, we analysed behavioural reports following awakenings (see Methods for details). The reported consciousness state varied across sleep stages as indicated by a chi-squared test (Χ^2^ = 26.3, p < 0.0001, Fig. 3a), with dreaming experience (DE) reports increasingly more frequent from N3 (32.1 % of awakenings) to N2 (56.2 %) and REM (87.9 %), in line with previous reports ^47^. No experience (NE) reports followed the opposite trend, decreasing from N3 (50 %), to N2 (21.9 %) and REM (0 %). We then checked whether the presence of experimental stimuli affected the report of a link between dream content and the presence of experimental stimuli before awakening (Fig. 3b), the frequency of reported consciousness state (Fig. 3c), and the type of sensory content experienced during dreams (Fig. 3d). A chi-Squared test showed no effect of stimulation on the frequency of the reported consciousness state (Χ^2^ = 0.136, p = 0.934). Sensory content ratings were not different depending on the presence of stimulation (all p-values > 0.21). Participants reported their experience to be related to the experimental stimuli in 33.3 % and 45.4% of trials, respectively, after stimulation and no stimulation blocks (Χ^2^ = 0.643, p = 0.42). Thus, there was no systematic incorporation of the multisensory stimuli used to probe PPS in dream content, and eventual reports of a link between the experimental stimuli and dream content are likely due to internally generated experiences (dreams about the experiment).

**Figure 3:**
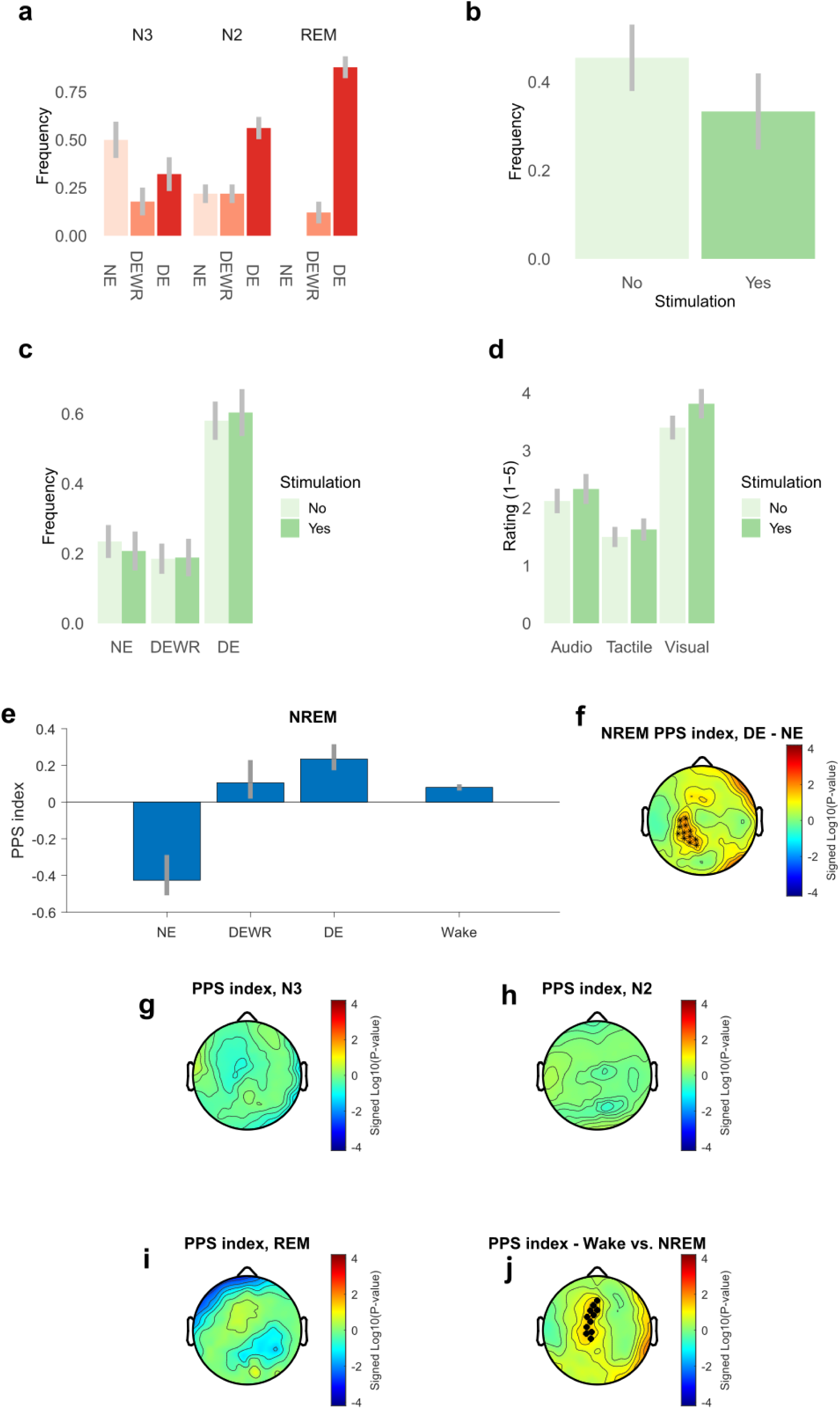
PPS index across sleep stages and reported consciousness states. (a) Frequency of NE, DEWR and DE reports depending on the sleep stage. (b) Frequency of reports of a link between sensory content of the dream and experimental stimulation, as a function of the presence of experimental stimulation. (c) Frequency of NE, DEWR and DE reports depending on the presence or absence of experimental stimulation before the awakening. (d) Average rating for the presence of Auditory, Tactile or Visual sensations depending on the presence or absence of stimulation. (e) PPS index in the high-beta cluster (identified during wakefulness) depending on the reported consciousness state at awakening, i.e., unconscious (NE), conscious without recall (DEWR) or conscious with recall (DE). The PPS index during wakefulness is also shown for comparison. Error bars show 66% confidence intervals obtained by bootstrapping. The analysis was conducted on NREM (N2+N3) trials. Panel (f) shows the contrast of PPS index in the DE vs. NE conditions. Asterisks indicate electrodes which are significant in this contrast and belong to the awake high-beta cluster. Panels (g), (h) and (i) show the PPS index in the N3, N2 and REM sleep phases, respectively. Panel (j) shows the contrast for the PPS index between wakefulness and NREM sleep. Black dots indicate electrodes belonging to the significant cluster.

Then, we moved to investigate our main hypothesis about the relation between PPS index and consciousness state. Following Siclari and colleagues ^37^, we analysed EEG responses during periods of 20 seconds immediately preceding an awakening in which stimuli were delivered. Due to the absence of NE reports in REM sleep, the analysis was only performed in NREM (N2+N3) sleep. We computed the high-beta PPS index for each category of trial in the clusters of electrodes identified during wakefulness, and, again following Siclari and colleagues ^37^, we contrasted DE and NE trials. The PPS index in DE was significantly higher than in unconscious (NE) states (p = 0.026), and not significantly different from wakefulness (p = 0.43). The PPS index in NE was also significantly lower than in wakefulness (p = 0.03). The effects were driven by both unisensory and multisensory stimuli presentations (see Fig. S1-S2). These differences could not be simply explained by an absence of evoked activity in NE trials, which exhibited robust responses compared to pre-stimulus baseline, both at the ERP and at the spectral level (see Fig. S3-S4). The PPS index in trials with dreaming experiences whose content could not be recalled (DEWR) and DE trials was not significantly different (p = 0.67), ruling out a possible confounding effect due to the ability of recalling dreams.

Next, we validated this finding beyond the pre-selected set of channels used in the previous analysis, focusing on the key contrast between NE and DE. We directly computed the DE – NE difference in PPS index and studied the resulting topography (Fig. 3f). The topography of electrodes resembled the one observed during wakefulness, with 9 out of the 29 channels within the high beta cluster identified during wakefulness showing a significant DE-NE difference.

Finally, we checked whether any significant PPS effect emerged in N3, N2 or REM sleep, irrespective of the reported consciousness state. No significant cluster of electrodes emerged in either phase (Fig. 3g-i). A significant cluster of electrodes emerged when contrasting and wakefulness and NREM sleep, for which we expect a lower PPS index (p = 0.035, one tailed, Fig. 3j). When directly contrasting REM and N3 sleep, respectively the sleep stage with the highest and the lowest percentage of DE reports, a trend for a higher PPS index in REM sleep emerged in a fronto-central channels (see Fig. S5). However, such trend did not survive cluster-based correction, possibly due to the limited number of trials available from the REM phase.

Taken together, these results show a neural marker of PPS representation emerged in conscious states, i.e., during wakefulness and dreams, and vanished during unconscious sleep (Fig. 3e), suggesting that the PPS index allows to identify the presence of consciousness during sleep.

### Spectral markers of PPS representation index consciousness levels and predict clinical outcome in patients with DoC behavioural phenotype

The above results show that our PPS index is a putative marker of consciousness in healthy participants. In patients exhibiting a DoC behavioural phenotype, the presence of residual consciousness, even when not detectable through routine bedside examination (covert consciousness ^48^), in the acute phase is possibly linked with higher chances of positive long-term outcome ^49^. We hypothesized that the PPS index could detect residual consciousness levels as assessed by standard clinical measures (i.e., the Coma Recovery Scale—Revised, CRS-R). In addition, we also expected that the PPS index, being independent from patients’ behavioural responses, could also detect covert consciousness, predicting patients’ outcome beyond standard clinical assessments.

To test this hypothesis, we adapted the experimental paradigm used in healthy participants to bedside recordings in clinical settings (see Methods, as in ^33^), and collected data from 72 patients in the acute neurorehabilitation unit. Recordings were performed on average 31.7 days (SD = 17.7) post brain injury. Based on the CRS-R evaluation at admission, patients were diagnosed with a DoC behavioral phenotype. Complementary evaluation using the motor behaviour tool (MBTr ^50,51^, see Methods) classified 67 out of 72 patients as clinical CMD (c-CMD), and the remaining 5 as true DoC (see Table S1 for clinical details and diagnoses according to CRS-R and MBTr). At discharge (23.5 days after the EEG recording, SD = 14.1), patients’ recovery was assessed by expert clinicians through a standardised examination that included four clinical scales evaluating motor, functional, and cognitive aspects. These scales were combined into a composite score providing a robust indicator of long-term outcome (outcome index, see Methods and ^52^). We investigated whether the PPS index at admission into the acute neurorehabilitation unit could predict the composite outcome measure at discharge.

Since the clinical 16-channels EEG setup covers centro-parietal, sensorimotor areas, largely corresponding to the cluster of high-beta modulation found in the healthy awake participants, we averaged the high-beta PPS index across the 16 channels. The high-beta desynchronisation PPS index correlated with patients’ level of consciousness as assessed clinically by the CRS-R at the time of the recording (R = 0.35, p = 0.002, Fig. 4a). Patients who were diagnosed as c-CMD also had a significantly higher PPS index (p = 0.019, Fig. 4c). These results confirm the link between PPS representation and consciousness levels in patients with DoC behavioural phenotype ^33^. Importantly, the PPS index was significantly and positively correlated with the outcome index (R = 0.43, p = 0.0001, Fig. 4b) measured at patients’ discharge: patients with more positive PPS index (i.e., in the direction of healthy awake participants) demonstrated a better recovery. Interestingly, the correlation of the PPS index with the outcome index, capturing patients’ state at discharge, was stronger than the correlation with the CRS-R, capturing patients’ behavioural responses at the time of the recording, even if the former was measured on average three weeks after the recording. These results were not driven by smaller or absent responses in patients with a worse outcome, since evoked responses to the stimulation in general were present irrespectively of the outcome (see Fig. S6).

**Figure 4.**
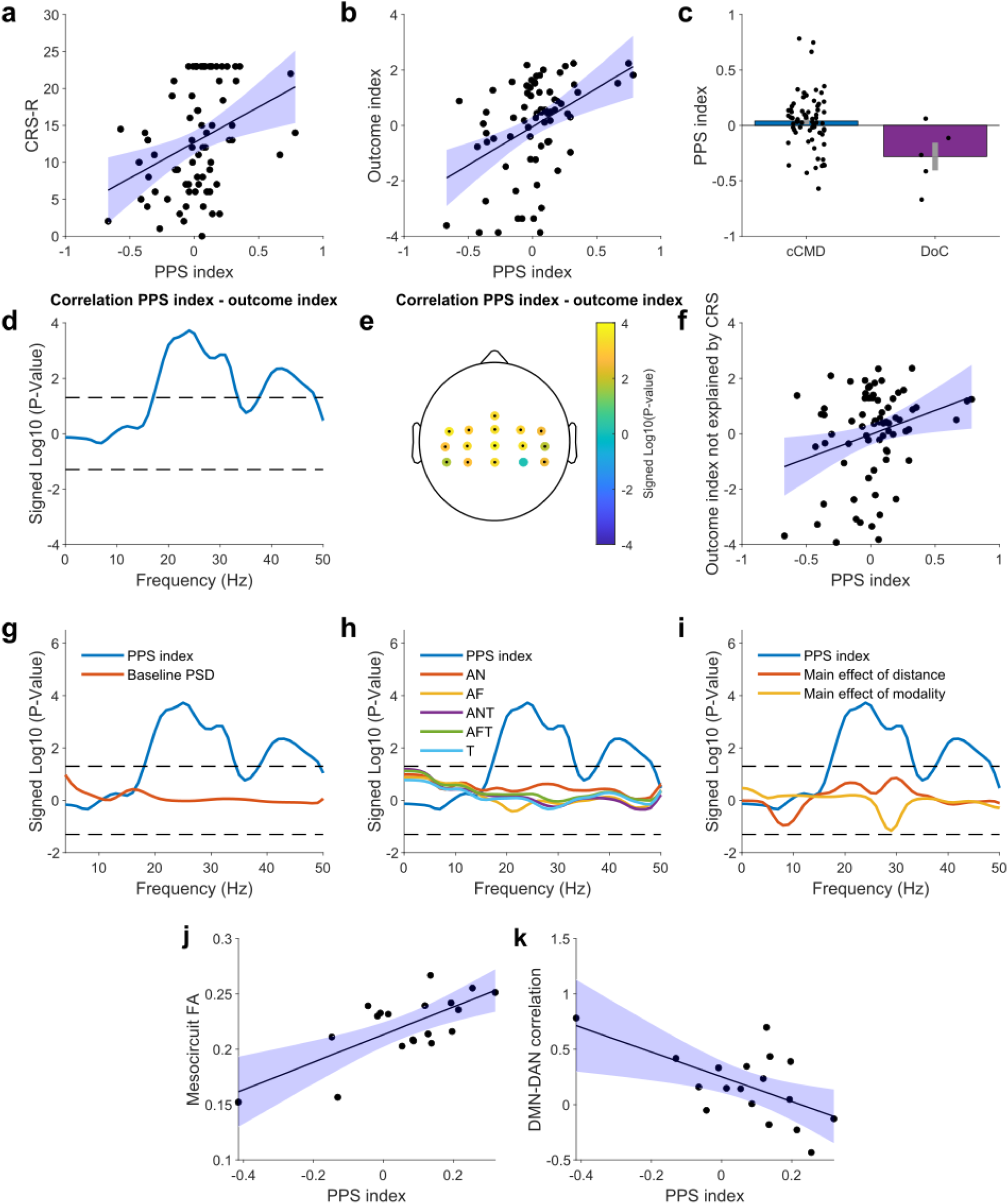
PPS index, consciousness and clinical outcome in patients with a DoC behavioural phenotype. (a) Correlation between PPS index (high beta, average on the 16 channels) and CRS-R at recording time in the 72 tested patients. (b) Correlation between PPS index and our composite clinical outcome index at discharge. (c) PPS index in patients who were diagnosed with c-CMD (left) and true DoC according to the MBTr. Error bars represent standard errors. (d) P-values for the correlation between PPS index (average on 16 channels) and outcome index at the frequency level. Positive values indicate a positive correlation, and vice versa for negative. Dashed lines indicate the significance threshold p = 0.05. (e) Correlation between high beta PPS index and outcome index at the single channel level for all the channels in the clinical EEG setup. Black dots indicate significant channels at p < 0.05. (f) Correlation between PPS index and outcome index, after regressing CRS-R scores at the time of the recording. (g-h-i) P-values from the same analysis as in panel b but comparing the PPS index with other potential predictors of clinical outcome, namely: (g) PPS index vs. power spectral density during the 0.5 s pre-stimulus baseline period; (h) PPS index vs. high beta power in all the individual experimental conditions (tactile T, audio-near AN, audio-far AF, audio-tactile near ANT, audio-tactile far AFT); (i) PPS index vs. the main effect of distance and modality. (j) Correlation between PPS index and forebrain mesocircuit integrity (fractional anisotropy, FA). (k) Correlation between PPS index and DMN-DAN connectivity in the same cohort of patients.

To investigate the specificity of these results, we then correlated the outcome index and the PPS index frequency by frequency. The PPS index significantly correlated with the outcome index in a cluster of frequencies spanning the 18-33 Hz frequency range (Fig. 4d), and peaking at 24 Hz, mainly within the high beta band, consistently with the results of the previous experiment. At the channel level, all but one channel showed a significant correlation between the high beta PPS index and the outcome index (Fig. 4e), with slightly stronger effects on the left side. This further supports the use of the average PPS index across all channels in this setup.

We then asked whether the PPS index could predict the clinical outcome beyond standard clinical information (CRS-R) available at the time of the recording. We thus tested the correlation between the PPS index and the outcome index while adding the CRS-R at the time of the recording as a covariate. The correlation between the PPS index and the outcome index remained significant (p = 0.021). To visualize this result, we regressed out from the outcome index the variability explained by the CRS-R. The residual variability in the outcome index not explained by CRS-R significantly correlated with the PPS index (R = 0.28, p = 0.017, Fig. 4f), suggesting that the CRS-R and PPS index provide complementary information about the clinical outcome.

We sought to determine whether simpler features of EEG response could better predict the clinical outcome. We first checked whether the outcome was predicted by the power spectrum in the pre-stimulus baseline period, which has been linked to recovery ^53–55^. No significant correlation emerged, and the PPS index in the high beta band was a stronger predictor than the baseline power spectra at all frequencies (Fig. 4g). Next, we tested whether the power spectrum in response to any of the individual stimulation conditions correlated with the outcome index, potentially driving the relationship between PPS and outcome. None of the five single experimental conditions showed a stronger correlation with the outcome index, compared to the high beta PPS index (Fig. 4h). Finally, as an even more stringent control, we tested whether the main effects of distance and modality (i.e., the overall ability to discriminate stimuli) correlated with the outcome index. The PPS index outperformed both measures in predicting the outcome index (Fig. 4i).

We then studied the relationship between PPS index and brain lesions characterizing DoC behavioural phenotype patients by studying predictors of coma recovery previously identified in a subset of 18 patients who also underwent magnetic resonance imaging (MRI) ^56^. We first examined a marker of structural integrity (fractional anisotropy) of the anterior forebrain mesocircuit, principally composed of frontal cortical regions, thalamus, and striatum, that has been shown to correlate with the severity of DoC and recovery potential ^57,58^. This index was positively correlated with the PPS index (R = 0.71, p = 0.0006, Fig. 4j), indicating that patients with greater structural integrity in the mesocircuit also had a higher PPS index.

At the functional lever, patients’ recovery has been associated with the anti-correlation between the activity of the default mode network (DMN) and the dorsal attention network (DAN) ^57^. Such anti-correlation is robustly present in healthy awake participants ^59^, and its absence characterizes the severity of DoC ^60,61^. The DMN-DAN anticorrelation has been further associated with the integrity of the mesocircuit ^57^. In the present sample, we found that the DMN-DAN correlation was significantly and negatively correlated with the PPS index (R = -0.59, p = 0.01, Fig. 4k), meaning that patients with a higher PPS index also had a stronger anti-correlation between DMN and DAN.

To understand whether alterations in the PPS index predicting patients’ outcomes were better explained by our structural or functional markers (which were highly correlated, R = 0.74), we ran a linear regression including both the forebrain mesocircuit integrity and the DMN-DAN correlation to predict the PPS index. The whole model was significant (R = 0.75, p = 0.004). The coefficient was significant for mesocircuit integrity (p = 0.031), but not for DMN-DAN (p = 0.66). This suggests that a lesion to the mesocircuit may affect both the PPS index and the DMN-DAN correlation.

## Discussion

We tested the hypothesis that PPS representation underlies a minimal form of self-consciousness, and therefore an EEG index of PPS processing (i.e., body-centered multisensory integration) could detect changes in the level of consciousness. In healthy awake participants, we found that PPS representation during wakefulness is well indexed by patterns of stimulus-induced changes in high beta (20-30 Hz) oscillatory power in a centro-parietal cluster of electrodes, that source analyses localized in typical fronto-parietal PPS regions ^19^. We thus took this index as an electrophysiological correlate of PPS representation. Next, we compared this PPS index across different states of consciousness during sleep, i.e., in dreamless sleep, when consciousness vanishes, and during dreams, when consciousness is present ^36^. During dreaming sleep, the PPS index was significantly higher than during dreamless sleep, and not different from wakefulness, as if PPS representation switched on during periods of conscious experience in sleep and off during unconscious sleep. In a second experiment in patients with a DoC behavioural phenotype, the same index of PPS representation was linked with consciousness levels at the time of recording, as measured by the CRS-R, and predicted their clinical outcome at discharge.

A link between PPS representation and minimal selfhood has been proposed theoretically (e.g., ^11^), and is supported by empirical studies ^27,62^ ^19^. Siclari and colleagues ^37^ demonstrated that the content of dreaming experience is associated with activations of cortical areas that underlie processing of the same information or modality during wakefulness. Since PPS mediates body-environment interactions from a first-person perspective ^14,63^, we predicted PPS representation would be active during dreaming. Interestingly, a recent survey found that most interactions in dreams occur within PPS from a first-person perspective ^64^. Here, we characterized a possible neurophysiological underpinning of these phenomenological reports by showing that an EEG marker of PPS representation characterizes dreaming sleep. To the best of our knowledge, these results provide the first neurophysiological evidence of multisensory integration in sleep, which had previously been explored only based on subjective reports ^65^. Furthermore, we linked body-centered multisensory responses, that is PPS representation, to the presence of conscious experience during sleep. The relationship between PPS representation and dream content appeared unrelated to the incorporation into dream experience of external stimuli. Indeed, despite the fact that auditory and tactile stimuli did evoke neural responses both in conscious and unconscious periods, participants rarely reported incorporation of these stimuli in their dreams (which are most often visual ^65^), and not systematically after blocks of stimulation. Thus, possibly some incorporation did happen, but it was very indirect, and leading no consistent reports. More importantly, we believe that the PPS index does not directly measure awareness *of external stimuli,* but rather whether the brain is in a state which supports conscious experience in general, PPS being linked to the subject of experience, not to its object. In this view, the experimental audio-tactile stimuli delivered in our paradigm could be used to probe the PPS system, and thus the presence of consciousness, even if they did not get incorporated into conscious experience. This is reminiscent of a prominent method measuring the complexity of the neural response to transcranial magnetic stimulation (TMS) as an index of consciousness ^66^. TMS pulses are carefully masked not to elicit any conscious percept, yet the neural response they evoke provides a robust proxy of the presence of conscious experience. Our approach is also complementary to other approaches on the neural correlates of consciousness, which measure electrophysiological processing of sensory stimuli as a proxy of awareness of an object of experience ^67^, or analyse spontaneous brain activity without external stimulation to infer the presence of conscious processing ^68,69^.

Data from DoC phenotype patients further support the relationship between PPS and consciousness. The PPS index correlated with CRS-R collected at recording, confirming the link between consciousness state as observed clinically and PPS, in line with our previous findings ^33^. More strikingly, patients with a positive PPS index at admission (more similar to healthy awake subjects), presented higher chances of a more favourable clinical outcome at discharge, while patients with a negative PPS index, (more similar to unconscious sleep), had a less favourable outcome. Our results suggest that a specific neurophysiological marker of PPS representation is linked to consciousness levels in both healthy subjects and patients, despite important differences in the physiology of deep sleep compared to DoC, first of all its reversibility. Previously, we demonstrated that supplementing standard evaluation with a thorough clinical assessment focused on observing spontaneous motor behaviour (MBTr), allows to distinguish a group of patients as exhibiting signs of interaction with their environment (defined as c-CMD ^50,51^). Patients with c-CMD also present distinct recovery trajectories, showing significantly better long-term outcomes ^70–72^. Accordingly, these patients presented a higher PPS index than patients with a true DoC. Thus, quantitative and accurate markers of covert consciousness could complement available clinical assessment, which suffers limitations ^73,74^ and can lead to patients’ mislabelling to an extent estimated as high as 40% ^75^.

Interestingly, the PPS index correlated more strongly with the outcome index at patient discharge than with the CRS-R at the time of the recording. This may depend on a generic higher sensitivity of the outcome index, based on extensive clinical assessments at discharge, as compared to the sole CRS-R. More intriguingly, this may relate to intrinsic limitations of the CRS-R scale, only partially predicting the long-term clinical outcome, as exemplified by its limitations in CMD patients ^72^. The CRS-R scale is based purely on the presence of behaviors indicative of consciousness, thus capturing only overt signs of consciousness. In contrast, our PPS index, not requiring any active response (non-task related), is also sensitive to covert consciousness. Thus, measuring patients’ residual multisensory integration in PPS may capture differences in patients’ preserved consciousness and potential of interaction, not detected by the CRS-R, and thus may further contribute to the outcome prediction. Importantly, the link between the PPS index and clinical outcome could not be reduced to the response to any specific experimental condition, nor to any combination of conditions other than the PPS index (see e.g. Fig. 4g). This supports our overarching hypothesis that a body-centered, multisensory representation of potential interactions between bodily and external stimuli, detects an important component of consciousness.

Beta oscillations, here used as a marker of PPS representation, are thought to underlie feedback signals in cortical networks ^76,77^, and have been related to multisensory ^78^ and sensorimotor ^79^ processing. Recent studies have specifically linked individual differences in beta-band synchronization and PPS representation ^80^, and stimulus induced beta-band desynchronization with the activation of the PPS system ^45^. The role of beta oscillations in sleep and altered states of consciousness is less known. A recent study using intracranial recordings showed that high-gamma responses to auditory stimuli (reflecting local spiking activity) in auditory cortices were comparable between wakefulness and sleep. However, beta-band desynchronisation followed auditory stimuli during wakefulness, but not during sleep ^81^. Since the combination of feedforward (gamma) and feedback (beta) signalling characterises perception during wakefulness, they interpret the absence of beta-band feedback signals as a distinctive feature of unconscious states. Our results may reflect a similar mechanism, specifically linked to the space-dependent processing of information that underlies PPS representation. Furthermore, patients with more negative outcome present a reduced DMN-DAN anticorrelation ^57^, which is known to be altered during deep sleep or anaesthesia ^60,82^ and is considered a key mechanism to mediate interactions with the environment ^60^. Here, DMN-DAN anticorrelation was associated with a higher PPS index, and with preserved forebrain mesocircuit integrity. This suggests that both the DMN-DAN anticorrelation and the PPS index may constitute two emerging faces of structural and functional integrity of key circuits for consciousness, whose functionality may break down either temporarily, as in deep sleep, or permanently, in DoC patients.

The present study has few limitations. First, patients’ clinical outcome is assessed at hospital discharge, with no follow-up, thus limiting the long-term predictive value of the PPS index. Mounting evidence shows both heterogeneity in recovery and delayed recovery after brain injury (both in DoC patients ^83^ and CMDs ^84^). Some of our patients might have shown more favourable progress if they had been evaluated after a longer period. Second, further efforts are still needed to pinpoint the exact neural mechanisms and structures underlying our index of PPS representation, and how it relates to physiological and pathological changes in consciousness.

To conclude, by focusing on the subject of the experience, i.e., the Bodily Self, we demonstrated a link between a specific neurophysiological marker of PPS representation and consciousness levels in healthy subjects and patients. A better understanding of the neural mechanisms of PPS representation could thus improve our understanding of the neural bases of consciousness, of how it fades during natural sleep-wakefulness cycles or following brain injury, adding a new perspective to the current approaches mainly focusing on the object of consciousness ^5^. Furthermore, PPS representation is highly plastic ^14,85^, as it likely emerges from statistical regularities during body-environment interactions ^17^. Thus, the present findings may suggest new paths to restore it in order to improve DoC patients’ outcome.

## Methods

### Sleep experiment - experimental procedure

#### Participants and general procedure

Fifteen healthy volunteers (age 22 ± 2 years, range 19-25 years, 8 females) were screened for psychiatric, neurological and sleep disorders. The sample size was determined based on previous studies about sleep modulation with external stimulations, where typically 15 participants were recruited ^86–88^. All volunteers had a good sleep quality as assessed by the Pittsburgh Sleep Quality Index (PSQI score < 5 ^89^), the Insomnia Severity Index (ISI; scores: < 10 ^90^). None of the participants had an extreme chronotype, as determined using the Morningness-Eveningness Questionnaire ^91^ and none of them suffered from excessive fatigue as assessed by the Fatigue Severity Scale (FSS score ≤ 36 ^92^) or sleepiness with the Epworth Sleepiness Scale (ESS; scores < 10 ^93^). The study was approved by the ethical committee of canton de Vaud, Switzerland, and was performed with the understanding and written consent of each subject. The experiments were performed in the Lausanne University Hospital sleep center.

Due to technical issues with EEG triggers, data recorded during sleep for two out of 15 recorded participants were not analysable. Hence, analyses on sleep data have been conducted on the remaining 13 participants. We verified that the main PPS index during wakefulness in the high beta range, shown in the previous paragraph, could be replicated in this subset of subjects (see Fig. S7). Furthermore, one subject did not present periods of REM sleep, limiting the sample for REM analyses to 12 participants.

#### Stimuli

Stimuli could be either auditory (near or far), tactile, or a combination of the two. This results in three possible unisensory and two multisensory: audio near (AN), audio far (AF), tactile (T), audio-tactile near (ANT), audio-tactile far (AFT). Auditory stimuli consisted in 100ms long bursts of white noise, generated by a PC using in Eprime 2 (Psychology Software Tools). Sounds were delivered through one of two commercial loudspeakers placed either 30 cm above subjects’ chest (near), or 130 cm above the chest (far). Tactile stimuli consisted in 5 ms electrical stimulation (Digitimer DS7AH) to the chest, 2 cm underneath the manubrium sternum.

To determine the intensity of auditory stimuli, we started from determining the auditory threshold. The minimal volume necessary for a sound to be perceived, was determined for each individual by using an up-down staircase procedure to detect auditory threshold. In this task, participants were lying in a bed with eyes closed and were told to detect sounds of different intensities. The experiment involved three blocks of 30 trials each. At the beginning of each block, the sound level was set at a value above threshold for all the participants (46 dB). Then, there was a reduction of 2 decibels when the subject’s response was positive (e.g., “I hear a sound”) and an increase of 2 decibels in stimulus level when the response was negative (e.g., “I do not hear anything”). We averaged volumes played during the last ten trials of each of the three blocks, and computed a grand average based on the three block averages, corresponding to the auditory threshold. Finally, we computed the value for the auditory stimulation to be used for the PPS (PPS auditory stimulation) as the mean individual detection threshold plus 40%. With this procedure, we made sure to adapt the auditory stimulation to each individual’s perception, and also to send an auditory stimulation at an unobtrusive volume during sleep (mean 51 dB, SD = 4.01).

The determination for the tactile value was based on the participant’s subjective perception of a similar intensity compared to the PPS auditory stimulation. Therefore, we administered a task, in which subjects were lying in bed with eyes closed. The individual PPS auditory stimulation value was sent and, immediately afterwards, a tactile stimulation was administered. The electrical stimulation was a single, constant voltage, rectangular monophasic pulse. The participants had to decide whether the tactile stimulation needed to be “more intense” or “less intense”, in order to be equal to the auditory stimulation in terms of perceived intensity. The task began with a tactile stimulation above threshold for all the participants (5mA), and there was a reduction of 0.5mA in intensity when the subject’s response was positive (e.g., “The tactile intensity should be higher to match the sound intensity to reach the same intensity as the sound”) and an increase of 0.5mA when the response was negative (e.g., “I would like less tactile intensity to reach the same intensity as the sound”). The PPS auditory stimulation was administered between each tactile stimulation, and the task ended once the participant decided that the tactile stimulation was equal, in terms of intensity, as the PPS auditory stimulation. This final value was then chosen as the tactile stimulation level to be used for the PPS experiment (mean 8.7mA, SD = 5.5mA). The values obtained with this procedure were in the same range as those employed in previous PPS studies ^33^.

#### Protocol - awake session

Before the sleep session, participants were lying in the bed with their eyes closed, and performed three blocks of PPS stimulation while we recorded EEG. Each block of the PPS experiment consisted of 110 trials: 22 unisensory tactile stimuli (T), 22 unisensory auditory stimuli at a near distance (AN), 22 unisensory auditory stimuli at far distance (AF), 22 audio-tactile stimuli near (ANT) and 22 audio-tactile stimuli far (AFT). The order of these trials was fully randomized within each block and the inter-trial interval consisted of 2.5-3 seconds (5 random steps of 100 milliseconds). Each block lasted approximately 7 minutes. No response was required from participants.

#### Protocol - sleep session

After recording the awake session, participants slept from approximately 11:30 pm to 6:30 am, while EEG data was continuously recorded. During the night, 5-minute blocks with the same randomized 5 types of stimuli as in the awake PPS session were alternated with 5-minute blocks with no stimulation. The blocks were administered during a stable sleep: N2, N3 or REM (N1 was avoided) and the order of these ones was randomized and administered regardless of the sleep stage. About 9 times per night (SD = 1.7), subjects were awakened and responded to a questionnaire about the presence of conscious experience and its content (see consciousness questionnaire paragraph). The questionnaire could either follow a block with or without stimulation in randomized order.

#### Consciousness questionnaire

During the night, subjects were awakened by a computerized alarm sound lasting 1.5 s, following which they were instructed, by interphone, to verbally answer a questionnaire used to assess sleep-related experiences ^47^. After ten seconds from the alarm, subjects were asked to report what had been going through their mind a moment before. Participants indicated whether they had a conscious experience and could remember its content (DE), had a conscious experience but could not recall the content (DEWR), or had not experienced anything (NE). In case they had experienced something and could recall the content (DE), they were asked to rate on an integer scale from 1 to 5, how much their sleep-related experiences included sensory experiences (i.e., tactile, auditory, visual), and how much these sensory experiences had been integrated into their dream. Additionally, they were asked whether their dream was related to the experimental stimuli (yes/no), and asked for a free description of the content of their dream. Following DEWR or NE reports we directly asked the last question, that is, to rate how much they felt asleep or awake before the alarm (1 to 5).

### Sleep experiment - EEG recording

A high-density EEG system (256 channels, Electrical Geodesics, Inc., Eugene, Oregon) with 500 Hz sampling rate was used. Four electrodes located near the eyes were used to monitor eye movements, and electrodes overlying the masseter muscles to record muscle tone.

### Sleep experiment - EEG preprocessing

The EEG signal was band-pass filtered between 0.5 and 45 Hz offline, and a notch filter at 50 Hz was applied to remove AC noise. Sleep scoring was performed over 30 s epochs according to standard criteria ^94^. Channels containing artifactual activity were visually identified and replaced by interpolation over nearby channels using spherical splines (NetStation, Electrical Geodesic). Ocular, muscular, and cardiac artifacts were eliminated through of Independent Component Analysis (ICA) using EEGLAB routines ^95^. ICA components presenting activity patterns and component maps typical of artifactual activity were removed through visual inspection ^96^. Epochs containing awakenings during the sleep session were excluded from EEG analyses. ICA computation and bad component rejection was performed separately for each analysed sleep stage (N2, N3, REM). Stimulus-locked epochs ranging from -1 second to +2 seconds relative to the stimulus onset were extracted from continuous EEG data collected. Each epoch underwent baseline correction by subtracting the mean value of the baseline period across the temporal dimension for each electrode. All subsequent EEG analyses were performed on average referenced data. Preprocessing for the wake session was identical to that for the sleep session, excluding the sleep scoring.

### Sleep experiment - Cortical Source Localization

To localize the PPS effect at the cortical level, source localization was performed on the preprocessed EEG signals using GeoSource software (Electrical Geodesics, Inc., Eugene, Oregon). The forward model was constructed based on individualized electrode geocoordinates, and the source space was constrained to a total of 2447 cortical voxels with a spatial resolution of 7 mm. The inverse solution was computed using the standardized low-resolution brain electromagnetic tomography (sLORETA) algorithm, combined with Tikhonov regularization (λ = 10^-2) to account for variations in the signal-to-noise ratio. The software provided the 3D (xyz) temporal evolutions of the 2447 cortical voxels. MATLAB was then employed to reconstruct the 1-D activity for each cortical voxel by projecting their xyz activity onto their principal orientation using singular value decomposition (SVD).

### DoC phenotype patients - experimental procedure

#### Participants and general procedure

In this second study, we examined the EEG correlates of the audio-tactile PPS paradigm, which had previously been evaluated in healthy participants during both wakefulness and sleep, in true DoC and putative CMD patients. The dataset comprised seventy-two patients (19 females, age = 48.9 ± 17.6 years, range = 19–84 years). Participants provided their informed consent to take part in the study, which was approved by the local ethic committee of the Vaud canton, Switzerland. Consciousness levels and diagnoses were determined by clinical neurologists and neuropsychologists using the Coma Recovery Scale-Revised (CRS-R).

Each patient underwent multiple testing sessions (average 2.5, range 1 to 10) across the DoC spectrum, with each session containing an average of 2.8 number of blocks (SD=0.5, range = 1 to 3). The number of sessions per patient was influenced by clinical requirements, primarily the patient’s availability for research and the duration of their stay at the acute neurorehabilitation unit, University Hospital of Lausanne (CHUV), Switzerland. Here, we analysed only data from the first session, which was available for the largest cohort of patients.

The experiment was conducted on average 30.7 days post-brain injury (SD = 14.1, range = 6 to 83 days). At discharge, which occurred 24 days after the EEG recording (SD = 14.1, range = 1 to 64 days), patient recovery was assessed by expert clinicians using a standardized evaluation, which included four different clinical scales combined to evaluate potential residual cognition and motor intent (as detailed in *DoC patients - clinical evaluation* Session).

#### Stimuli

Auditory stimuli were presented at two different distances. Specifically, the stimulation consisted of 50 ms of white noise played through loudspeakers (Z120 Portable Speakers, Logitech, Lausanne, Switzerland) positioned at 5 cm and 75 cm (65.2 dB SPL and 64.1 dB SPL respectively) from the participant’s extended arm in the depth dimension. Tactile stimuli were equally delivered to the participant’s arm for 50 ms, at a frequency of 35 Hz, via functional electrical stimulation (FES; MEDEL Medical Electronics, MOTIONSTIM 8, Innsbruck, Austria) placing two electrodes (positive and negative, Flextrode Plus) on the extensor digitorum communis (dorsal part of the arm near the elbow). The stimulation intensity was set at 70% of each participant’s motor threshold, determined immediately before the experiment and ranging from 5 mA to 11 mA. In addition to the electrodes on the participant’s right arm, a third sensor was positioned on the right shoulder, serving as an earth ground to mitigate potential electrical artifacts from the FES stimulation. In the case of multisensory trials, auditory and tactile stimuli were administered synchronously.

#### Protocol

During the experiment, patients were positioned supine at an angle of approximately 130° within a controlled environment with regulated lighting and sound. Each block of audio-tactile stimulation lasted 10 minutes and comprised 250 trials, including 50 unisensory tactile stimuli, 50 unisensory auditory stimuli presented at a near distance, 50 unisensory auditory stimuli presented at a far distance, 50 audio-tactile stimuli at a near distance, and 50 audio-tactile stimuli at a far distance. The sequence of these stimuli was fully randomized within each block, with an inter-trial interval uniformly distributed between 1.5 and 2 seconds. Due to clinical considerations, the number of blocks recorded per patient varied, averaging - blocks per session and ranging from - to - blocks per session. Brief breaks were provided between blocks.

### DoC phenotype patients - clinical evaluation

Before their admission to the unit, patients were repeatedly evaluated clinically by neurologists and neuropsychologists using the Coma Recovery Scale-Revised (CRS-R) and the Motor Behavior Tool (MBTr ^50^). The MBTr is a clinical motor observation tool designed to describe the spontaneous (without stimulation) motor behaviour in a non-task related way (akinesia versus reflex motor pattern of decortication and decerebration) and detect subtle non-reflexive movements and “positive” responses that suggest intentionality, that are not considered by the CRS-R alone, along with a systematic search for obstacles limiting their interaction capabilities. Preserved conscious integration is considered present according to this tool when at least one positive item is noted, leading to the classification of the patient as having clinical CMD (c-CMD). The MBTr has demonstrated the ability to identify a subgroup of patients exhibiting motor intention, the presence of which predicts favorable recovery ^50,72,97^.

### DoC phenotype patients - EEG recording

Patients’ EEG data were collected using a 16-channel EEG system (g.USBamp and g.Nautilus, g.tec medical engineering, GmbH, Graz, Austria) with a sampling rate of 500/512 Hz and referenced to the right earlobe. Electrodes were positioned to cover motor and somatosensory areas (Fz, FC3, FC1, FCz, FC2, FC4, C3, C1, Cz, C2, C4, CP3, CP1, CPz, CP2, and CP4).

### DoC phenotype patients - EEG preprocessing

EEG signals were band-pass filtered offline between 0.5 and 40 Hz, with an additional notch filter to eliminate power line noise. Epochs were extracted from continuous EEG data spanning -100 ms to 500 ms relative to stimulus onset and were baseline-corrected to the 100 ms pre stimulus onset. Bad channels were excluded based on an amplitude threshold of ±100 µV, and bad epochs were rejected based on joint probability measures to ensure data integrity. Then, within each session, blocks of epochs were concatenated prior to performing Independent Component Analysis (ICA). Components with less than a 60% likelihood of representing neural activity were excluded from further analysis. Following this pre-processing procedure, six patients were excluded due to excessive noise and artifacts that compromised the EEG signals, resulting in a total of 175 sessions available for analysis. Finally, the EEG data were re-referenced to the average reference to standardize the signals for subsequent analyses.

### Sleep experiment - Spectral analyses

For each epoch, the Power Spectral Density (PSD) was computed using Welch’s periodogram method with a Hamming window of 0.5 seconds, 50% overlap between segments, and a frequency resolution of 1 Hz. PSDs were calculated separately for the baseline and post-stimulus. Subsequently, the PSDs of both baseline and post-stimulus were averaged across trials for each stimulus condition. The post-stimulus PSDs were then normalized relative to an average baseline PSD, computed as the mean PSD across all five conditions. This procedure resulted in a normalized post-stimulus PSD for each subject and stimulus condition (tactile, auditory near, auditory far, auditory near + tactile, and auditory far + tactile). Finally, the normalized post-stimulus PSDs were averaged across specific frequency bands: theta (4-8 Hz), alpha (8-12 Hz), low beta (12-20 Hz), high beta (20-30 Hz), and gamma (30-40 Hz).

### DoC experiment - Spectral analyses

For each epoch, the Power Spectral Density (PSD) was calculated for post-stimulus using Welch’s periodogram method with a Hamming window of 0.25 seconds, 50% overlap between segments, and a frequency resolution of 1 Hz. Subsequently, the PSDs were averaged across trials for each stimulus condition. The post-stimulus PSDs were then normalized relative to an average baseline PSD, computed as the mean PSD across all five conditions. This procedure resulted in a normalized post-stimulus PSD for each subject and stimulus condition (tactile, auditory near, auditory far, auditory near + tactile, and auditory far + tactile). Subsequently, after demonstrating that the average PSD across the 16 channels exhibits a PPS effect within the frequency band identified in the previous dataset, we focused exclusively on computing the spectral analysis in this frequency band (high-beta: 20-30 Hz). All subsequent statistical analyses of correlation with clinical scales were based on Pearson correlations.

### MRI indexes

Methods used to derive the MRI indexes described in Fig. 4 are presented in detail in ^56^. Dffusion weighted imaging (DWI) data was denoised, preprocessed, and used to derive fractional anisotropy (FA) maps as described previously in ^52^ We assessed the structural connectivity of the forebrain mesocircuit using the multi-scale probabilistic atlas of the human connectome ^98^. To calculate the mesocircuit structural connectome, we extracted the mean fractional anisotropy (FA) values from the voxels within the white matter bundles that connect the bilateral regions of the forebrain mesocircuit, including the frontal cortex, precuneus, cingulate cortex, thalamic nuclei, and basal ganglia. The biomarker for structural connectivity (mesocircuit FA) was then derived by averaging the FA values across the entire mesocircuit connectome.

To determine the DMN-DAN correlation we first performed an independent component analysis (ICA) to identify these brain networks. Anatomical and resting state fMRI data were preprocessed through the default fmriprep pipeline (21.0.2) ^99^. To compute resting-state functional connectivity we used data-driven, group ICA ^100^ to decompose the preprocessed and smoothed data in 20 spatially ICs. We selected the ICs into the components corresponding to resting state networks, discarding noise components through visual inspection and comparison to the resting state networks templates ^101^. The mean group spatial t-value maps of the remaining ICs were thresholded at t > 4 and used as brain masks for further analyses. The DMN-DAN correlation was then defined as the Pearson correlation of the signal time course in the corresponding masks.

### Statistical analyses

#### PPS index - definition and statistical testing

Most previous electrophysiological studies ^33,34^ on PPS representation analysed signals in the time domain, comparing event related potentials (ERPs) or global field power (GFP) across different stimulation conditions. Here, we applied the same approach to frequency domain features of the response to stimuli, which also show modulations related to PPS processing ^45,80,102^. Indeed, transient evoked activity reflected by ERPs is known to change broadly in amplitude, latency and topography between wakefulness and sleep ^103,104^ or in disorders of consciousness ^105–107^. Thus, to derive a PPS-based marker that can be reliably compared across conscious states, here we developed a novel EEG measure of PPS representation based on the oscillatory correlates of stimulus processing. Similarly to what was done in previous electrophysiology studies, the PPS index was defined based on the key functional property of PPS representation, that is, a multisensory and body centered representation linking external stimuli (in this case, auditory) and body-related stimuli (tactile). In terms of comparisons between experimental conditions, this translates to searching for electrodes demonstrating a stronger near-far difference for multisensory stimuli than for unisensory ones, leading to the formula

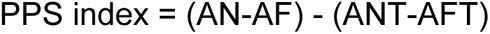

This analysis was performed on the normalized power averaged on the 1 second window following each stimulus for each frequency band. For convenience, since lower power in the key high beta band indicates higher neural activation, we reversed the sign of the formula with respect to studies focusing on evoked potentials, where higher amplitude corresponds to higher activation. This way, the PPS index is expected to be positive in case of high beta power.

#### Cluster-based statistics

To assess the statistical significance of differences in Power Spectral Densities (PSDs) across channels, we employed a cluster-based permutation test ^108^ focusing on predefined frequency bands. This analysis was conducted using a custom MATLAB script integrating functions from the FieldTrip toolbox. We specifically analyzed 187 internal electrodes, excluding more peripheral ones such as those on the cheeks. The Monte Carlo method was employed to determine statistical significance, involving 5000 random permutations of observed values between two conditions. Specifically, we applied a two-tailed permutation-based t-test for dependent samples (alpha = 0.05) and implemented cluster-based correction on the spatial dimension. The test statistic was evaluated using the maximum sum under the permutation distribution. To account for spatial adjacency between channels, we computed a neighbour structure for the internal electrodes. Clusters were defined as groups of at least four significant neighbouring channels to ensure robustness in spatial clustering.

#### Source Level Statistics

To evaluate significant voxel activations in the high-beta PPS index, a two-tailed permutation-based t-test for dependent samples against zero was conducted, utilizing 5,000 random permutations and setting a threshold at p < 0.05 (uncorrected) for displaying the results.

#### Awakening questionnaires

These analyses aimed at establishing a link between PPS index and reported consciousness state (NE, DEWR, DE). The analyses were performed on awakenings following PPS stimulation blocks, and restrained to NREM sleep due to the limited number of trials in NREM sleep. Similarly to what done in a previous work on the neural correlates of dreaming, we analysed responses to stimuli occurring in the 20 seconds preceding each awakening. Not all subjects presented at least one awakening for each of the three possible consciousness reports, nor a full set of the four experimental conditions needed to compute the PPS index. Thus, we could not simply average PPS indexes across subjects and consciousness reports and perform regular parametric statistics. To overcome this, we used non-parametric statistics based on bootstrapping to estimate the variability of our data and compare consciousness reports. We randomly resampled (with replacement) 13 participants from our original data and computed the PPS index by pooling all trials from all subjects for each consciousness report. The values obtained through this procedure should mimic the distribution of hypothetical replications of the experiment, allowing to estimate confidence intervals on PPS index values and p-values ^109^. Confidence intervals (shown in Fig. 3e) were estimated as the central 66% of the distribution of bootstrapped PPS index values. P-values were computed by measuring the proportion of resamples yielding larger (or smaller) values in condition A than in condition B. In total, 62 trials in the NE condition, 37 trials in the DEWR condition, and 104 trials in the DE condition were analysed.

### Linear Mixed Models

To investigate how the PPS index interacts with the state of consciousness during sleep, we employed linear mixed models (LMMs). Our objective was to determine whether variations in stimulus distance (near vs. far) and modality (unisensory vs. multisensory), along with the presence (DE) or absence (NE) of dreaming, influence EEG power before the awakenings. For this analysis, the power in beta-high frequency band (20-30 Hz) was considered, since the PPS effect was previously observed in this range during wakefulness. To analyze the three-way interaction, we applied linear mixed models separately for each channel using the following formulation: *power ∼ distance * modality * dreaming + (1|subject)*. In this model, *distance*modality* captures the PPS effect, while the term *dreaming* indicates the presence (DE) or absence (NE) of dreaming. The model included random intercepts for each subject to account for individual variability in beta-high power.

## Supplementary Materials

**Figure S1.**
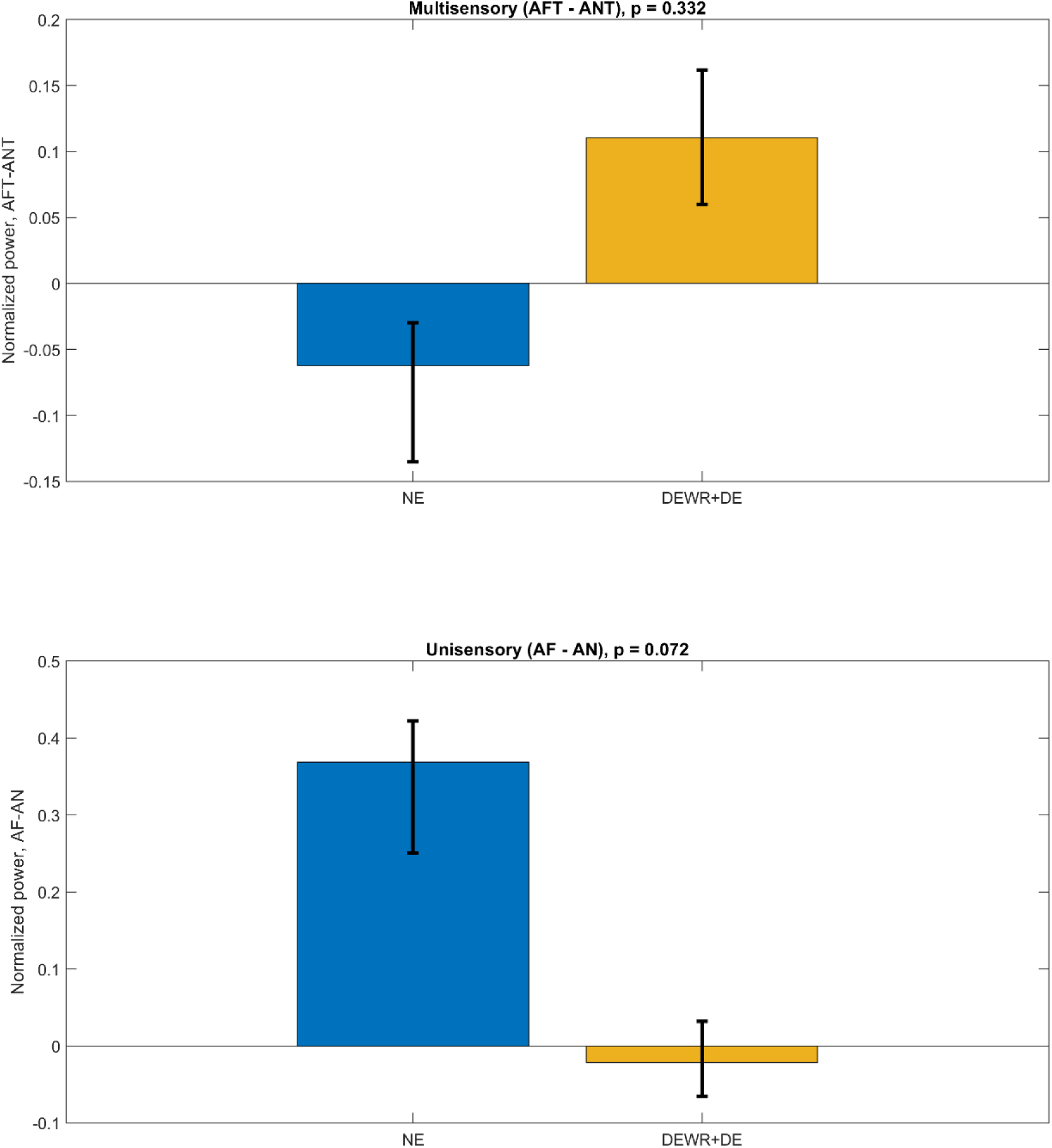
Near-far difference in normalized power for multisensory (top) and unisensory (bottom) trials, compared between unconscious (NE) and conscious (DEWR+DE) sleeping conditions. The p-value refers to the conscious-unconscious comparison.

**Figure S2.**
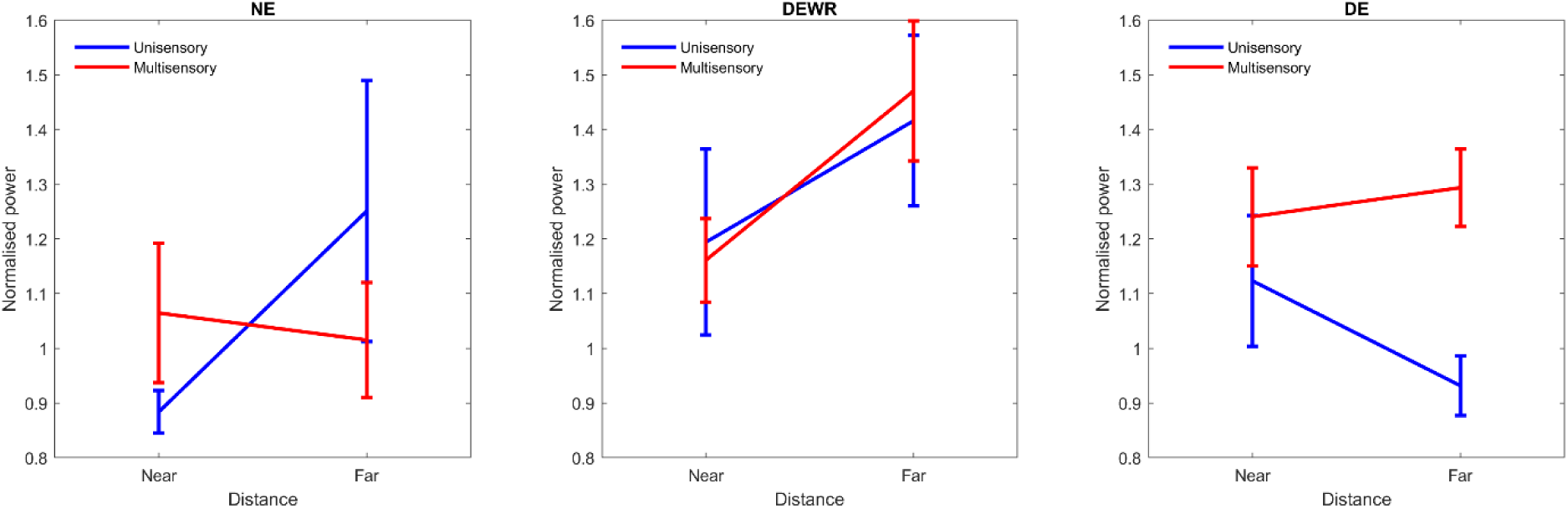
Comparison of normalised high-beta power between consciousness levels showing all the four experimental conditions contributing to the PPS index.

**Figure S3.**
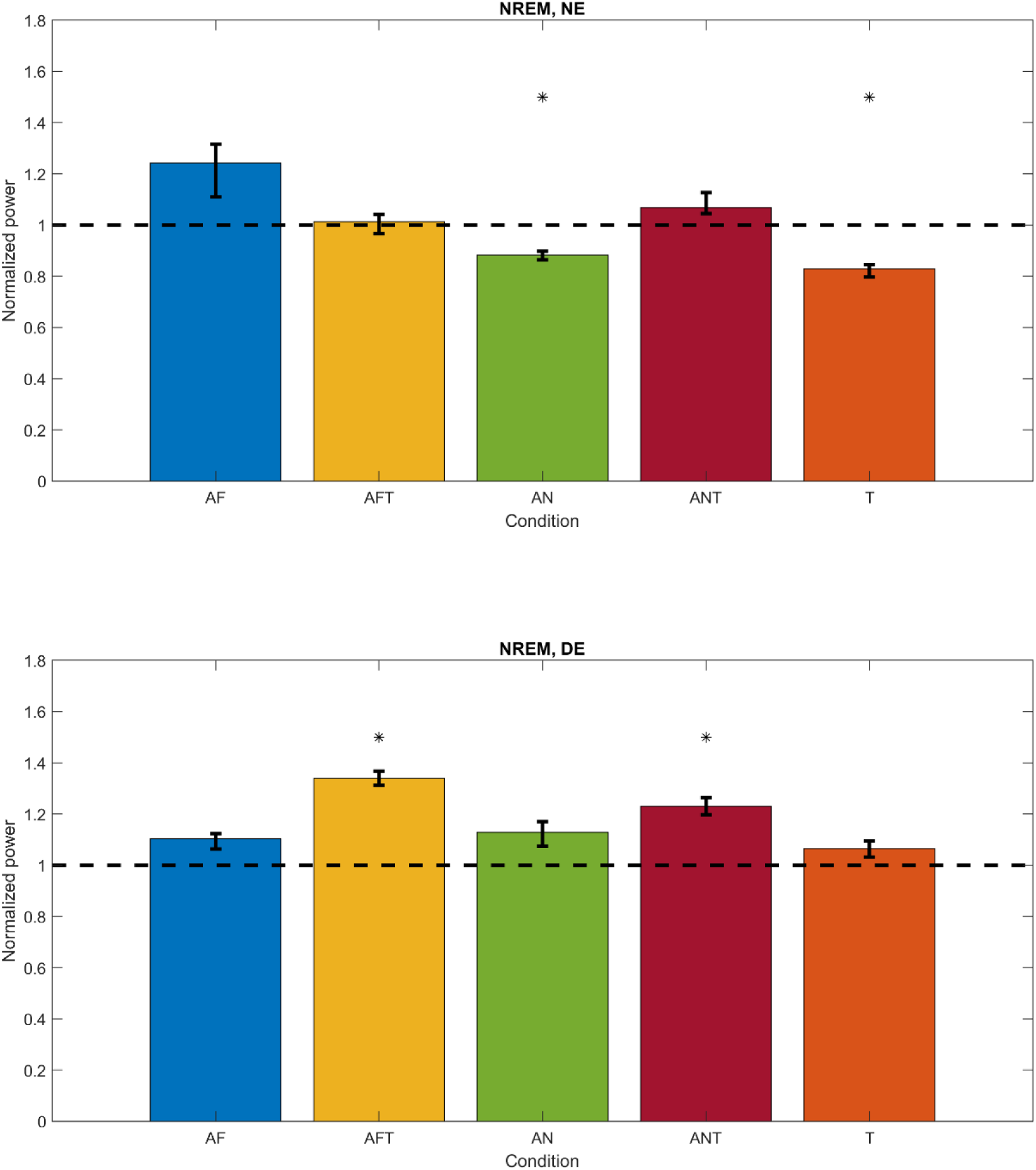
Comparison between stimulus induced response and baseline for the five experimental conditions, in NE (top) and DE (bottom) trials. Asterisks denote conditions showing a significant difference from baseline (the 1 second pre-stimulus window used for normalization).

**Figure S4.**
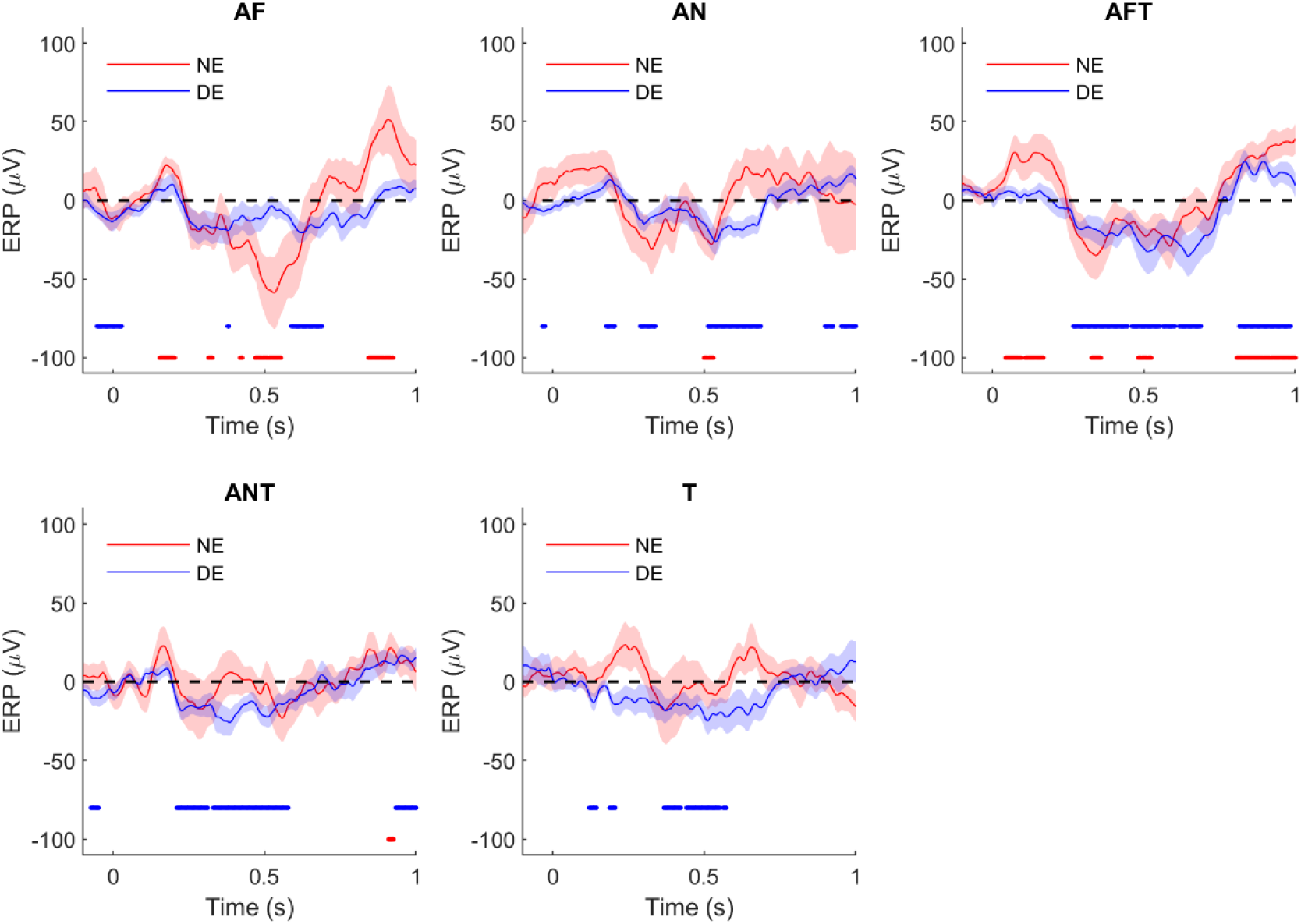
Comparison between stimulus induced response and baseline for the five experimental conditions, in NE (red) and DE (blue) trials. Dots below the plots denote timepoints showing a significant difference from 0 for NE.

**Figure S5:**
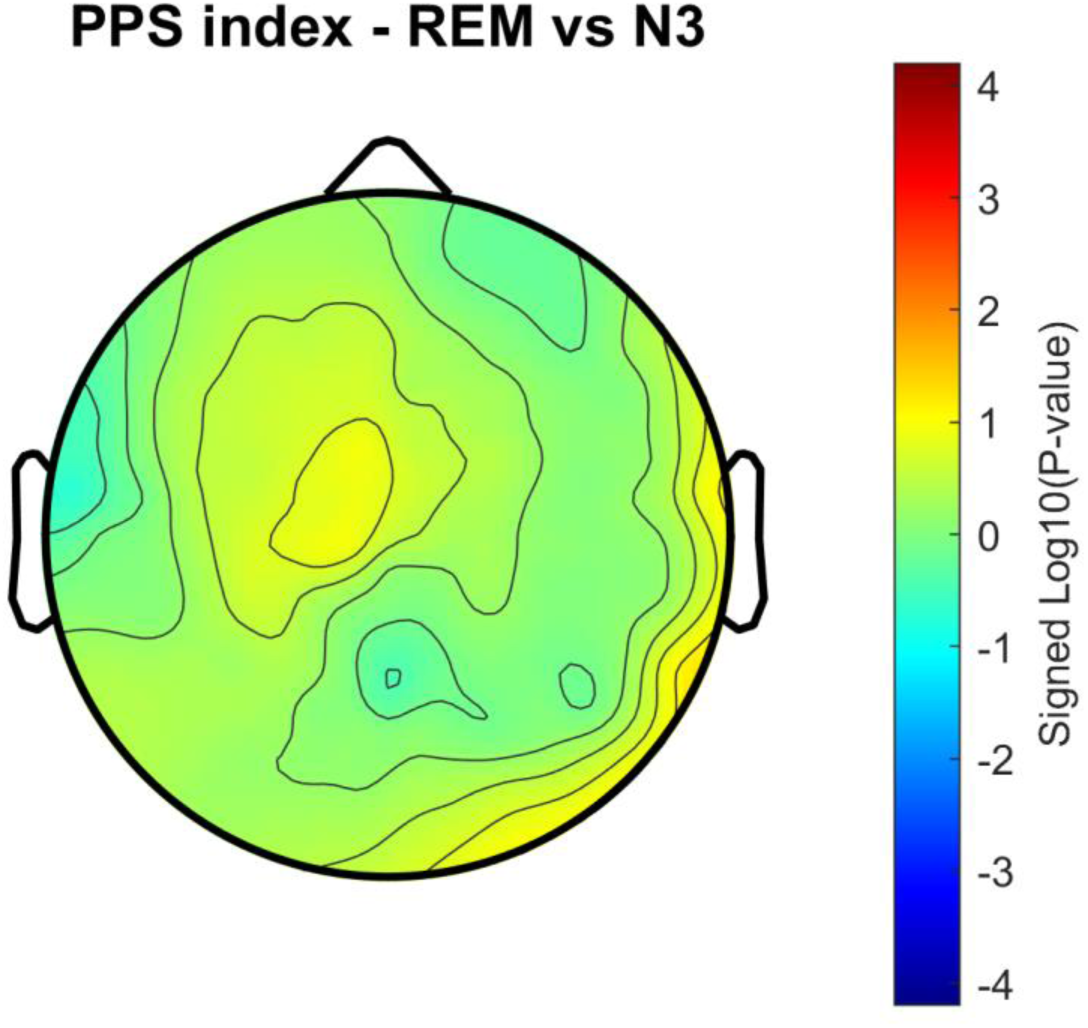
contrast between REM and N3 PPS index. Positive values indicate a higher PPS index for REM compared to N3.

**Figure S6.**
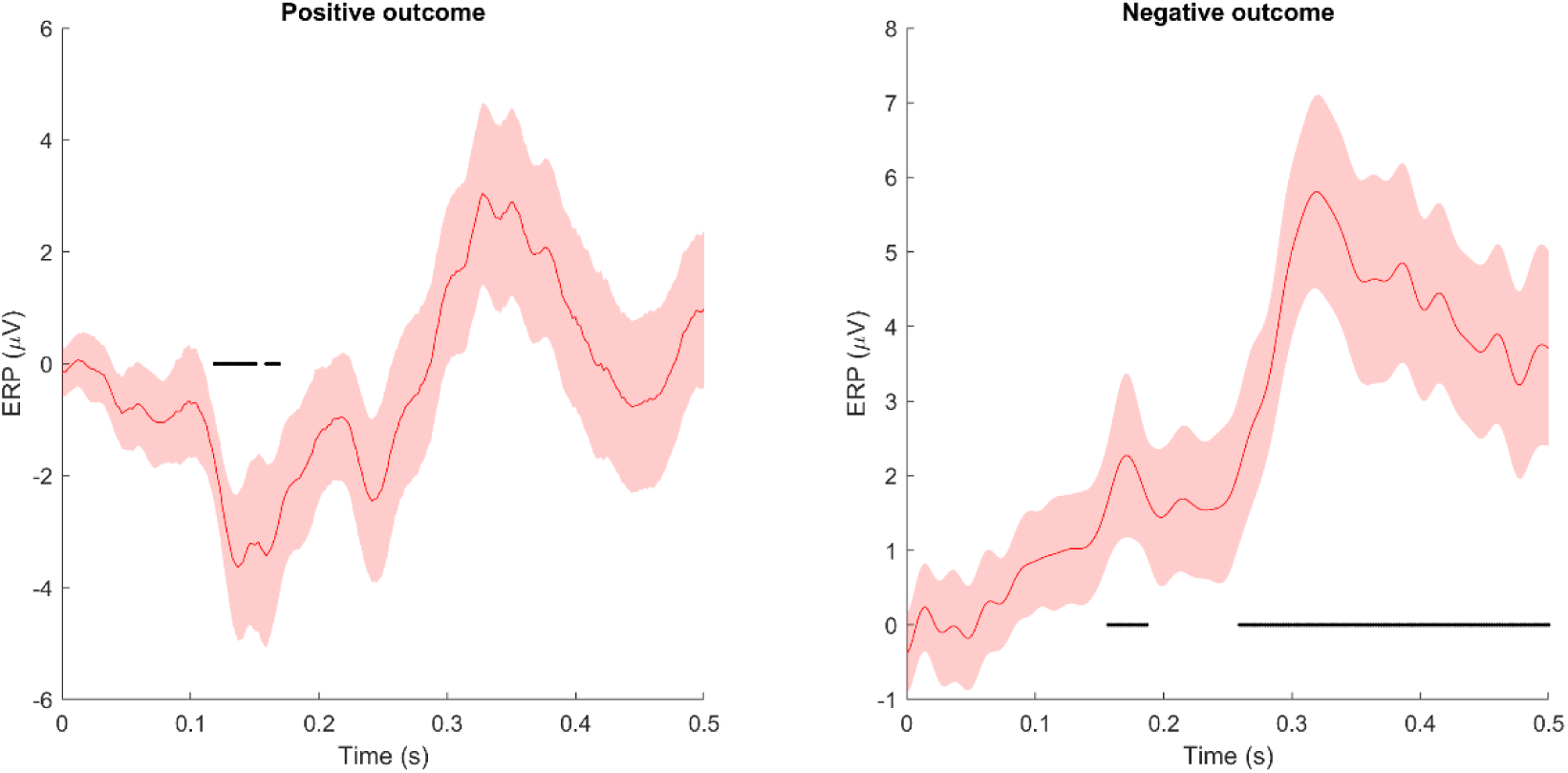
Example of response (sum of AN, ANT, AF, AFT) from the C3 channel in positive and negative outcome patients (based on a median split). Shades denote standard errors, black lines indicate significant timepoints compared to the pre-stimulus baseline.

**Figure S7:**
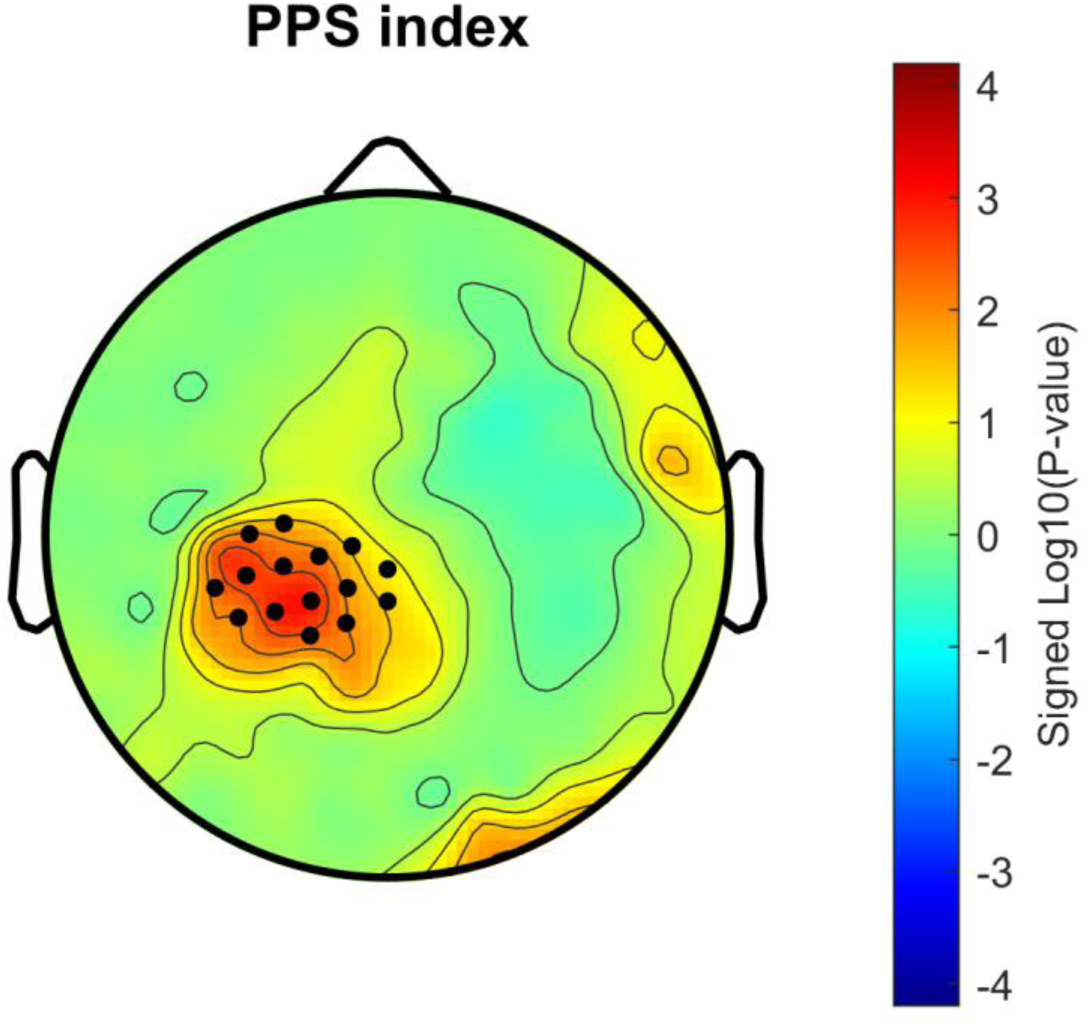
PPS index as in figure 2a, but computed only for the subset of 13 participants with analysable sleep data.

**Table S1.**
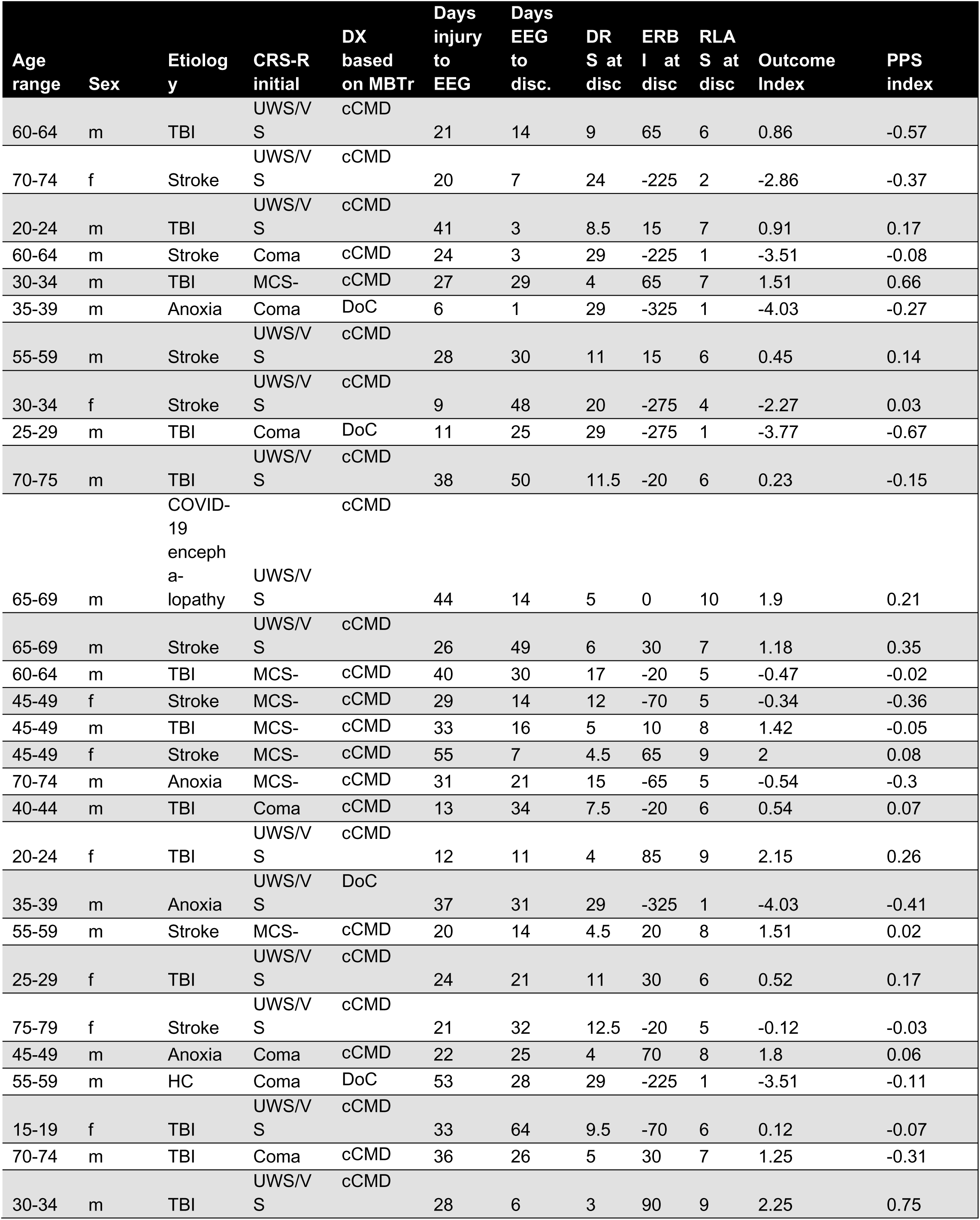
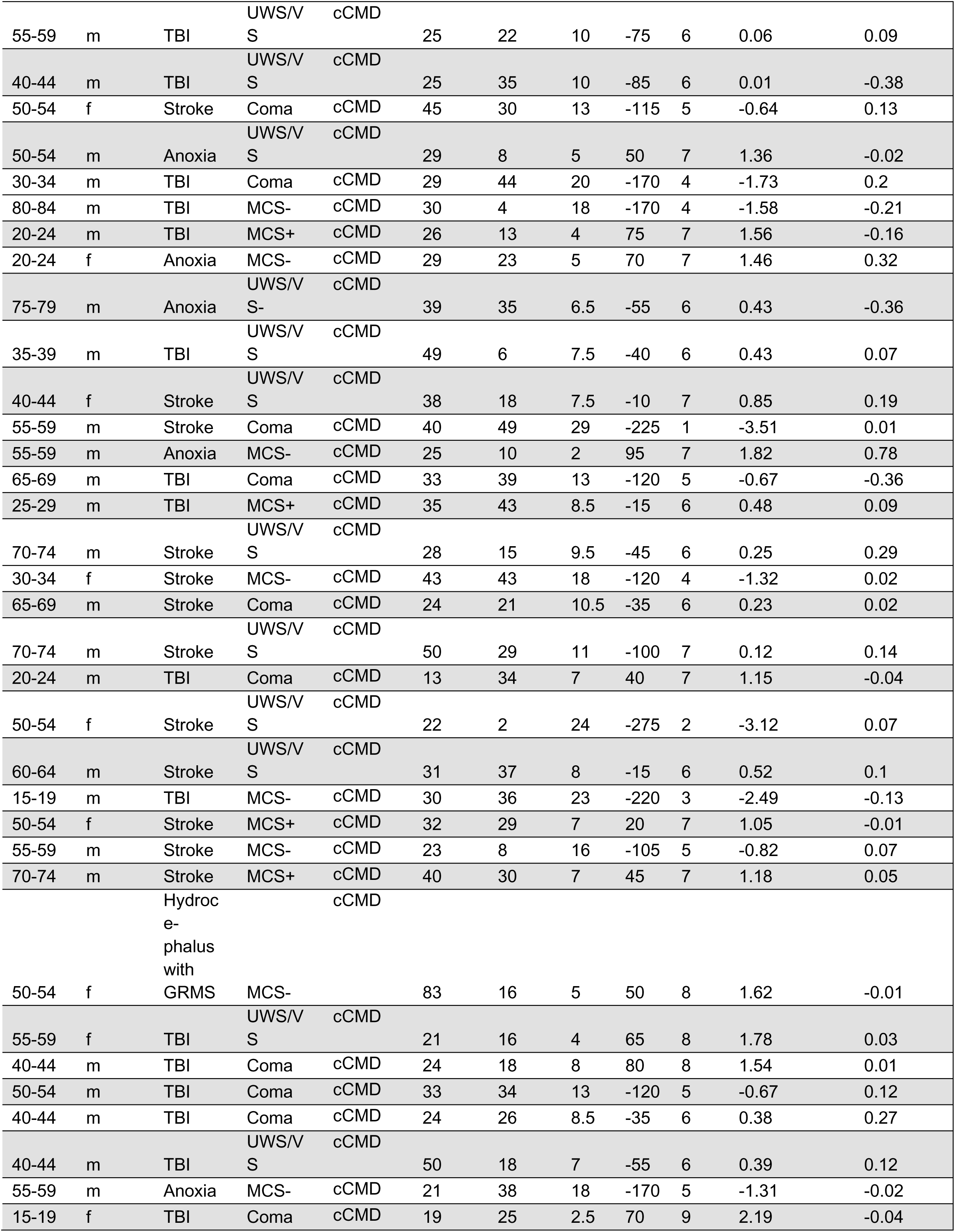
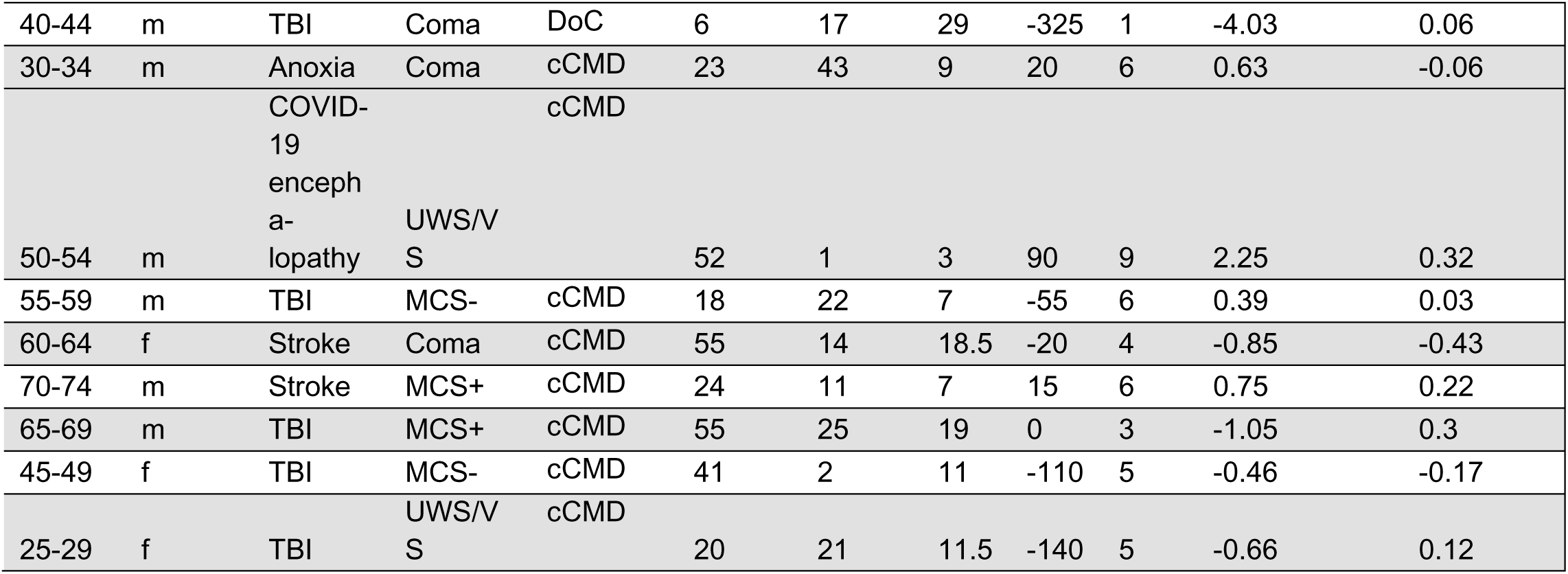
Demographic and clinical data. TBI traumatic brain injury, GRMS Global Rostral Midbrain Syndrome, CRS-R Coma Recovery Scale Revised, UWS/VS unresponsive wakefulness syndrome/vegetative state, MCS-/+ minimally conscious state minus/plus, EMCS emergence of minimal conscious state, MBTr Motor Behavior Tool revised, cCMD clinical cognitive motor dissociation, DRS Disability Rating Scale, ERBI Early Barthel Index, RLAS Rancho Los Amigos Level of Cognitive Functioning. Category on the Early Barthel Index range from -325 to 100, with 100 indicating complete functional independence; Category on the Disability Rating Scale range from 0 to 29, with 0 indicating absence of disability; Category on the Rancho Los Amigos Levels of Cognitive Functioning range from 1 to 10, with 10 indicating modified independent. Good outcomes were considered as ERBI ≥ -75, DRS < 11, RLAS ≥ 6.

